# Stochastic Epigenetic Dynamics of Gene Switching

**DOI:** 10.1101/2020.03.18.996819

**Authors:** Bhaswati Bhattacharyya, Jin Wang, Masaki Sasai

## Abstract

Epigenetic modifications of histones crucially affect the eukaryotic gene activity. We theoretically analyze the dynamical effects of histone modifications on gene switching by using the Doi-Peliti operator formalism of chemical reaction kinetics. The calculated probability flux in self-regulating genes shows a distinct circular flow around basins in the landscape of the gene state distribution, giving rise to hysteresis in gene switching. In contrast to the general belief that the change in the amount of transcription factor (TF) precedes the histone state change, the flux drives histones to be modified prior to the change in the amount of TF in the self-regulating circuits. The flux-landscape analyses elucidate the nonlinear nonequilibrium mechanism of epigenetic gene switching.

---

Stochastic nonlinear nonequilibrium dynamics of complex systems have been analyzed by the combined description of probability flux and landscape, where the landscape represents the distribution of states generated via nonlinear interactions and the flux reflects the nonequilibrium driving force of transitions among states [1–3]. In particular, the combined flux-landscape approach provided a transparent view of the coupled dynamics of continuous and discrete variables such as the interplay between concentration changes and transitions of biomolecules [4, 5]. In this Letter, we study a cell biophysical problem of coupled continuous and discrete dynamics, the epigenetic dynamics of gene regulation, by developing a method of flux-landscape analyses.

Epigenetic pattern formation associated with chemical modifications of histones provides long-term memory of gene regulation, which plays a critical role in cell differentiation, reprogramming, and oncogenesis [6]. The effects of epigenetic modifications have been discussed theoretically [7–16]; however, their quantitative dynamics still remain elusive. Statistical mechanical analyses have suggested that an array of ∼10^2^ interacting nucleosomes (i.e., particles of histone-DNA complex) show collective changes in their modification pattern [17–19]; such collective modifications have been experimentally observed by comparing the chromatin structural and histone modification data [20–22], and the dynamics of collective histone modifications have been observed in cells with an engineered promoter site [23, 24]. Thus, the theoretical and experimental analyses highlighted the dynamical role of collective histone modifications in gene switching.

Similar to in the previous models [8, 17, 18], we classify the collective histone states around a promoter site of a modeled gene system into three states: the gene-activating state with histones marked by acetylation as H3K9ac or others (*s* = 1), the gene-repressing state as the state with histones methylated as H3K9me3 or others (*s* = −1), and the neutral state with mixed or an absence of activating and repressing histone marks (*s* = 0). The chromatin chain with *s* = 1 takes an open structure, which facilitates access of the large-sized molecules for transcription to DNA to enhance the gene activity, whereas the *s* = –1 chromatin is condensed, which prevents the access of necessary molecules to suppress the gene activity (Fig. 1). We write the transition rate from state *s*′ to *s* as 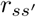. We assume that the transcriptional activity of the gene is regulated by both the histone state and binding of a transcription factor (TF); TF binds to DNA near the promoter site (*σ* = 1) or unbinds from DNA (*σ* = −1) with binding rate *h* and unbinding rate *f*. These rates should be insensitive to the chromatin packing density when the size of TF is as small as the pioneer TFs which can diffuse into the compact chromatin space [25, 26]. Here, for simplicity, we consider that the TF is a pioneer factor and its binding/unbinding rates are not affected by the chromatin compactness or the histone state. However, the bound TF should recruit histone modifier enzymes as observed in engineered cells [23, 24], which modifies 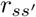 as 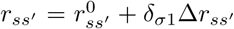, where *δ*_*σ*1_ is a Kronecker delta. The rate of collective change in many histones, 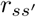, should be smaller in general than the rate of single-molecule binding/unbinding, *h* or *f*, which enables histones to retain the effect of TF binding as memory; 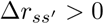 for *s* > *s*′ when the TF works as an activator while 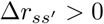 for *s* < *s*′ when the TF works as a repressor.

**FIG. 1.**
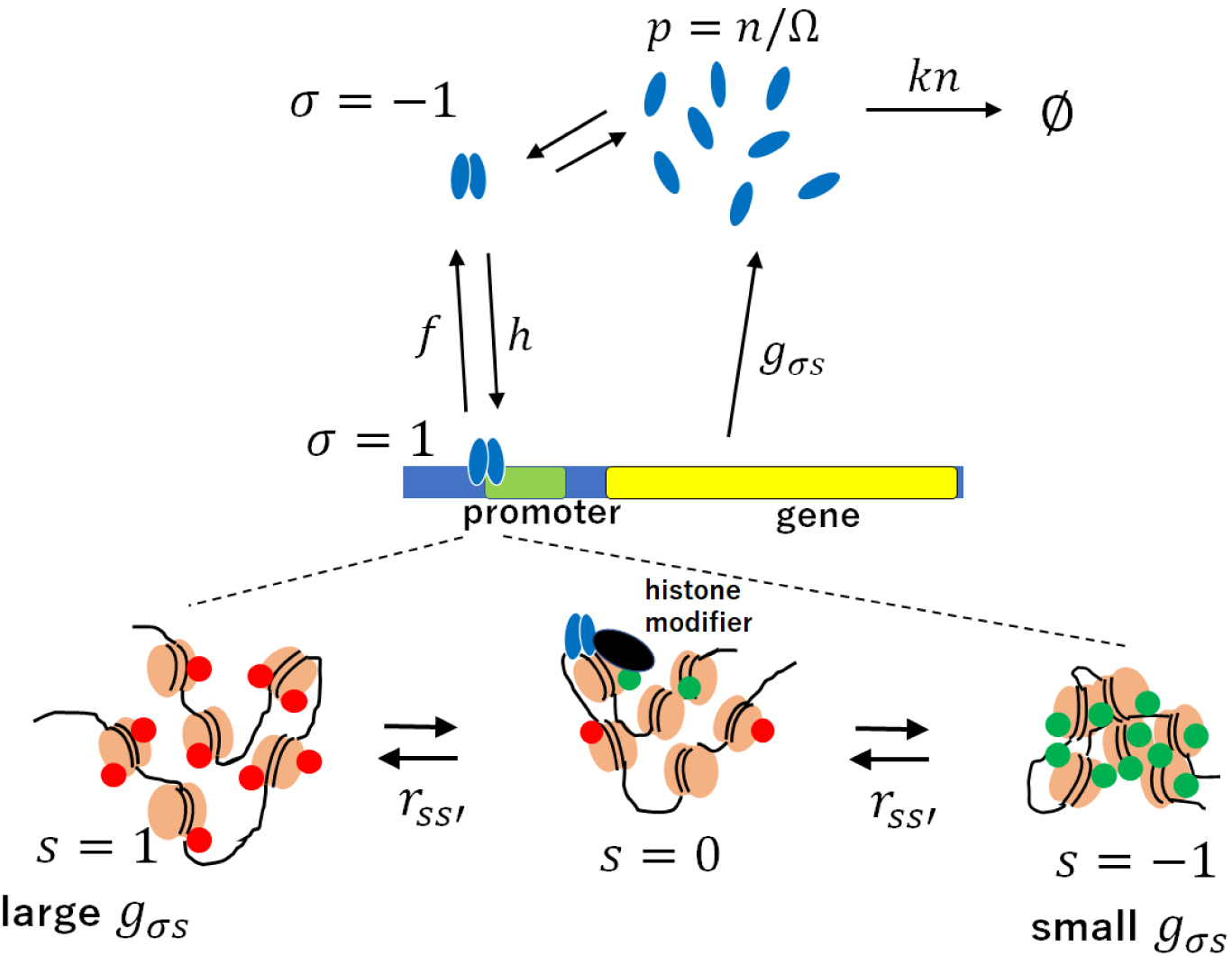
Schematic of a self-regulating gene. Close-up views of chromatin at around the promoter region are illustrated at the bottom, where histones (brown) are actively marked (red) or repressively marked (green). Histones in the chromatin are collectively marked through their mutual interactions. When the activating mark is dominant, chromatin has an open structure, which enhances the gene transcription activity (*s* = 1), and when the repressive mark is dominant, condensed chromatin suppresses the gene activity (*s* = −1). When neither mark is dominant, the hisotne state is neutral (*s* = 0). The transcription factor (TF) is a dimer of the product protein (blue oval). The bound TF recruits histone modifiers (black), which perturb the transition rates 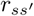 among the histone states. The protein production rate *g*_*σs*_ depends both on the TF binding status, *σ*, and on the histone state, *s*. See the text for the definition of rates denoted on the arrows.

As a prototypical motif of nonlinear gene circuits in cells, we consider a self-regulating gene as illustrated in Fig. 1; a dimer of the gene product protein is the TF acting on the gene itself. Here, by assuming that the dimerization reaction is much faster than other reactions, the TF binding rate can be written as *h* = *h*_0_*p*^2^, where *p* = *n/*Ω is the product protein concentration, *n* is the protein copy number in the cell nucleus, Ω is the typical copy number in a volume of ∼ *µ*m^3^ in the nucleus, and *h*_0_ is a constant. Through transcriptional regulation by TF binding and histone modifications, the protein production rate depends on both *σ* and *s* as *g*_*σs*_. Here, *g*_11_ is the largest and *g*_−1−1_ is the smallest of *g*_*σs*_ ≥ 0 for an activator TF, whereas *g*_−11_ is largest and *g*_1−1_ is smallest for a repressor TF. The protein degradation rate is *kn* with a rate constant *k*. Thus, the histone state is a part of the feedback loop of gene regulation, constituting a relatively slowly varying part in the loop.

To describe the reactions in Fig. 1, we use the operator formalism of Doi and Peliti [27–29], which has been applied to problems of gene regulation without explicitly considering the epigenetic degree of freedom *s* [4, 30–33] and to the problem of histone state fluctuation without considering the TF binding status *σ* [19]. Now, taking into account both *σ* and *s*, we consider the probability that the protein copy number is *n* at time *t, P*_*σs*_(*n, t*). We define a six dimensional vector ***ψ***(*t*), whose component is ***ψ***(*t*)_*σs*_ = Σ _*n*_ *P*_*σs*_(*n, t*)|*n* >. The operators *a* and *a*^†^ are introduced as *a*^†^|*n* >= |*n* + 1 >, *a*|*n* >= *n*|*n* − 1 >, and [*a, a*^†^] = 1. Then, by assuming that all the reactions in Fig. 1 are Markovian processes, the master equation of the reactions is 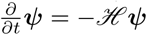, with *ℋ* being a six dimensional operator,

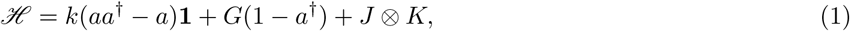

where **1** is a unit matrix, *G* is a diagonal matrix, 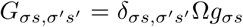, and *J* and *K* are transition matrices for *σ* and *s*, respectively;

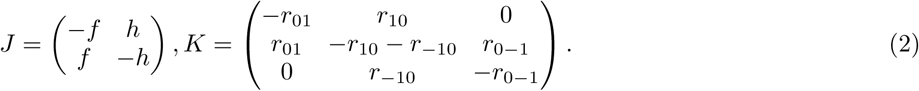

Here, the nonequilibrium features of the reactions are reflected in the non-Hermiticity of *ℋ*.

The temporal development of ***ψ*** can be represented in a path-integral form by using the coherent-state representation, 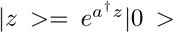, with a complex variable *z*(*t*). The transition paths in the *σ* and *s* space can be represented by using the spin 1/2 and spin 1 coherent states with spin angles *θ*(*t*) and *α*(*t*) and their conjugate variables *ϕ*(*t*) and 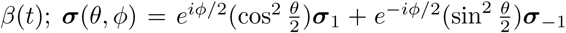 with ***σ***_*i*_ = (*δ*_*i*1_, *δ*_*i*−1_)^*T*^ for TF binding/unbinding and 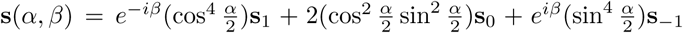 with **s**_*i*_ = (*δ*_*i*1_, *δ*_*i*0_, *δ*_*i*−1_)^*T*^ for the epigenetic histone degree of freedom. Then, from the saddle-point approximation of the path integral of variables *z, θ, ϕ, α*, and *β*, we obtain the Langevin equations describing fluctuations in the protein concentration *p*, the TF binding status *ξ* = cos *θ*, and the histone state *ζ* = cos *α*;

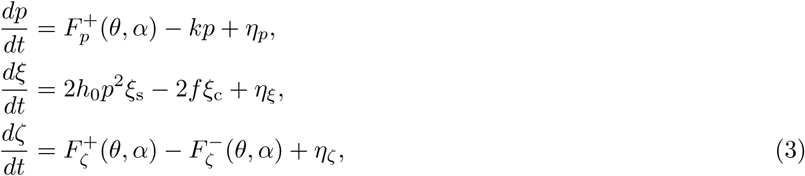

where

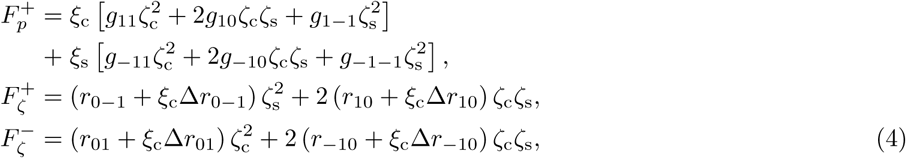

with *ξ*_c_ = cos^2^(*θ/*2), *ξ*_s_ = sin^2^(*θ/*2), *ζ*_c_ = cos^2^(*α/*2), and *ζ*_s_ = sin^2^(*α/*2). In Eq. 3, *η*_*p*_, *η*_*ξ*_, and *η*_*ζ*_ are mutually independent Gaussian white noise with an average of 0 and dispersions *B*_*p*_, *B*_*ξ*_, and *B*_*ζ*_;

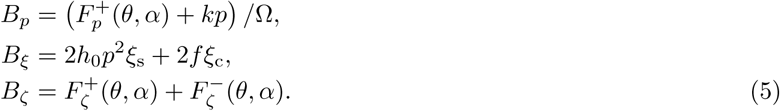

We should note that *p* = *n/*Ω was an almost continuous variable for a large Ω. Hence, the original coupled dynamics of a nearly continuous variable *p* and the discrete variables, *σ* and *s*, were transformed into the Brownian dynamics in the 3-dimensional (3D) space of the continuous variables, *p, ξ*, and *ζ*, with TF bound (*ξ* = 1)/unbound (*ξ* = –1) and histone state activating (*ζ* = 1)/neutral (*ζ* = 0)/repressing (*ζ* = –1).

In a single-molecule tracking experiment of the TF in embryonic stem cells (ESCs), the observed time scale of binding/unbinding was ∼ 1/min [26], whereas the observed degradation rate *k* of TF was ∼ 0.1/hour [34, 35]. In engineered ESCs, the collective histone state was changed with the rate of ∼ 1/day [23]. Therefore, the estimated ratios are *h/k* ∼ *f/k* = *O*(10^2^) and 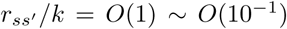. With such large *h* and *f*, TF binding/unbinding reactions can be treated as *adiabatic*: TF binding/unbinding are regarded as in quasi-equilibrium as ⟨*dξ/dt*⟩ = 0, leading to *ξ*_s_ = 1/ [(*h*_0_*/f*)*p*^2^ + 1]. With this adiabatic approximation, the 3D calculation in Eq. 3 for (*p, ξ, ζ*) is reduced to the 2D calculation for (*p, ζ*).

Either with adiabatic or non-adiabatic treatment of TF binding/unbinding kinetics, the flux-landscape approach [1, 3–5] provides a concise view of the results obtained by numerically integrating Eq. 3. With the adiabatic approximation, for example, the landscape, *U*(*p, ζ*), is obtained from the calculated stationary probability density distribution as *U*(*p, ζ*) = − ln *P* (*p, ζ*). The probability flux **J** is obtained as a 2D vector field in the adiabatic case from the Fokker-Planck equation corresponding to Eq. 3, 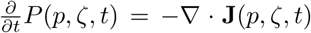, where ∇ = (*∂*_*p*_, *∂*_*ζ*_) and **J** = (*J*_*p*_, *J*_*ζ*_) with 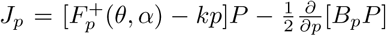 and 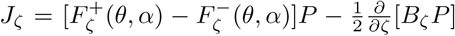. Of note, even in the stationary state, **J** can be nonzero when ∇· **J** = 0. This divergence-less circulating flux is a hallmark of the breaking of the detailed balance and reflects the hysteresis in switching dynamics [2–4].

Fig. 2 shows the calculated *U* and **J** at the stationary state in the adiabatic case. For an activator TF, when the TF binding affinity is small (small *h*_0_*/f*) (Fig. 2a), *U* has a single basin at *p* ≈ 0 and *ζ* ≈ − 1 ∼ 0 (off state). When the binding affinity is intermediate (intermediate *h*_0_*/f*) (Fig. 2b), *U* has two coexisting basins in the off state and at *p* ≈ 0.7 and *ζ* ≈ 1 (on state). When the binding affinity is large (large *h*_0_*/f*) (Fig. 2c), a dominant basin is found at the on state. Thus, the binding affinity of the activator TF determines the distribution of stable states and works as a switch that determines both the histone state and gene activity in a correlated manner. When the TF is a repressor, we find a single basin at intermediate *p* and *ζ* for a wide range of binding affinity (Fig. 2d).

**FIG. 2.**
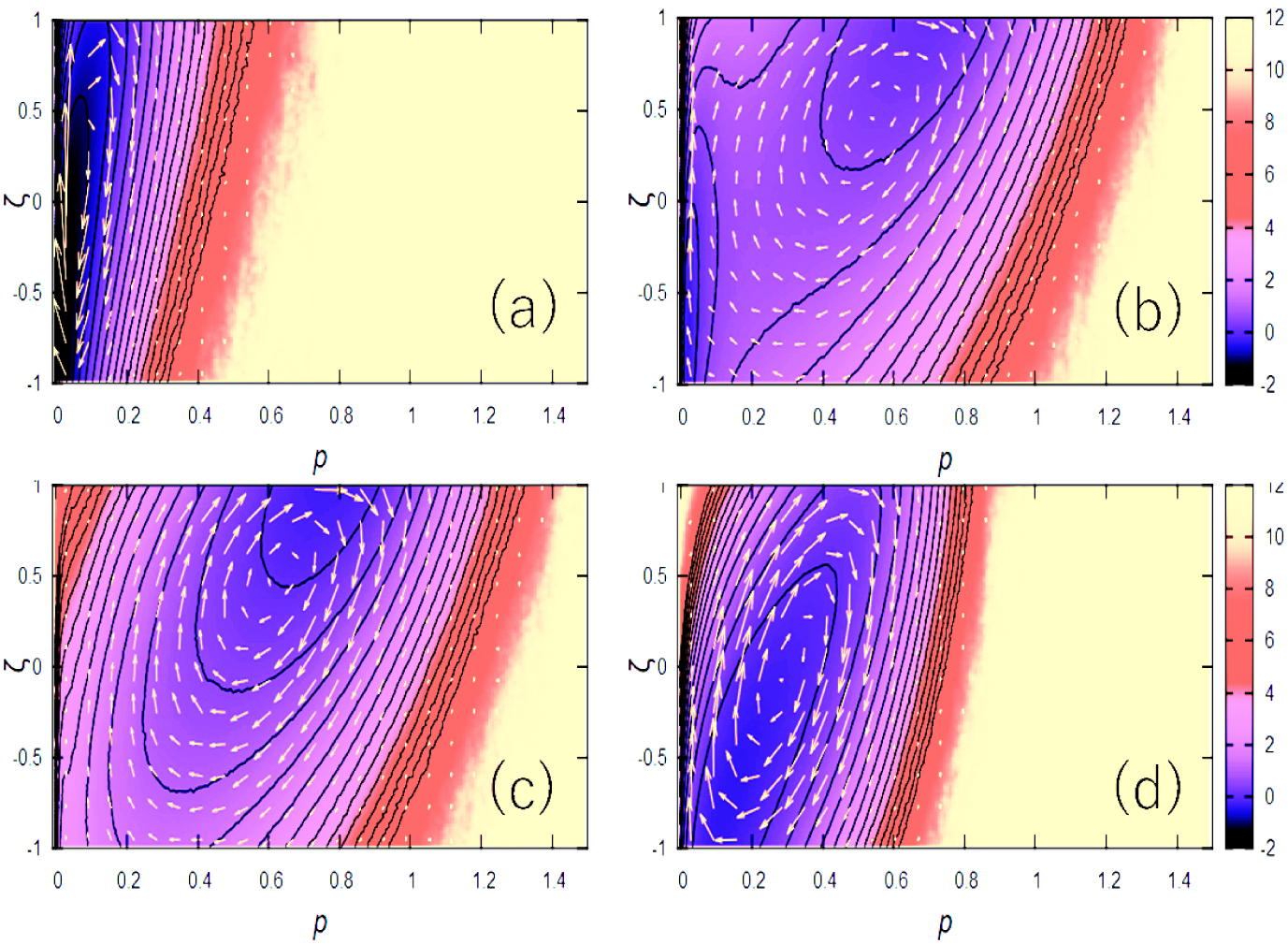
Landscape *U*(*p, ζ*) and probability flux **J**(*p, ζ*) calculated on the 2D plane of the protein concentration *p* and the histone state *ζ* in the adiabatic approximation of TF binding/unbinding. *U* is shown with contours and **J** with yellow arrows. (a–c) Self-activating and (d) self-repressing genes. (a) *h*_0_*/f* = 2, (b) *h*_0_*/f* = 20, (c) *h*_0_*/f* = 200, and (d) *h*_0_*/f* = 2. The rate parameters are scaled in units of *k* by setting *k* = 1; *g*_11_ = 1, *g*_−11_ = *g*_10_ = 0.2, *g*_−10_ = *g*_1−1_ = *g*_−1−1_ = 0, Δ*r*_10_ = Δ*r*_0−1_ = 1, Δ*r*_01_ = Δ*r*_−10_ = 0 for (a)–(c) and *g*_−11_ = 1, *g*_11_ = *g*_−10_ = 0.2, *g*_10_ = *g*_1−1_ = *g*_−1−1_ = 0, Δ*r*_10_ = Δ*r*_0−1_ = 0, and Δ*r*_01_ = Δ*r*_−10_ = 1 for (d). For (a)–(d), Ω = 100 and *r*_10_ = *r*_0−1_ = *r*_01_ = *r*_−10_ = 1.

In all the cases shown in Fig. 2, we find a flux **J** globally circulating around a basin or traversing between basins. When the epigenetic effect is absent, the flux is diminished in the adiabatic limit [4, 5], but here with non-adiabatic epigenetic histone dynamics, the circular flux is significant even in the limit of adiabatic TF binding/unbinding. The evident circulating flux suggests a temporal correlation between the histone modification and the gene activity change. By following the flux direction along the path from the off to the on states, the histone state first tends to become activating and then the protein production follows, and from the on state to the off state, the histone state first tends to become repressive, and then, protein production follows.

This temporal correlation is confirmed by calculating the normalized difference between the two-time correlation functions, *A*(*t*) = [⟨*δζ*(*τ*)*δp*(*τ* + *t*)⟩ – ⟨*δp*(*τ*)*δζ*(*τ* + *t*)⟩] /| ⟨*δζ*(*τ*)*δp*(*τ*)⟩|, with *δζ* = *ζ –* ⟨*ζ*⟩ and *δp* = *p* − ⟨*p*⟩, where ⟨…⟩ is the average over *τ* and over the ensemble of calculated trajectories. We can write *A*(*t*) ∝⟨det [**q**(*τ*), **q**(*τ* + *t*)]⟩ with a 2D vector **q**(*τ*) = (*ζ*(*τ*), *p*(*τ*)). A positive value of *A*(*t*) at *t* > 0 implies that the increase (decrease) in *ζ* tends to increase (decrease) *p* at a later time *t*. For activator TF (Fig. 3a) and repressor TF (Fig. 3b), *A*(*t*) is plotted for various values of 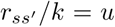, showing that *A*(*t*) has a peak with a positive value at *t*_*u*_ = 1*/u ∼* 0.5*/u*. We find that the peak is evident even in the case when the histone dynamics is as slow as *u* < 1, which indicates that the prior change in the histone state to the gene activity is not owing to the faster rate of reactions in histones but is due to the circulating flux generated by the non-adiabatic slow dynamics of histones. The delayed influence of *ζ* on *p* should induce hysteresis in the switching dynamics; *ζ* tends to decrease prior to the decrease in *p*, while *ζ* tends to increase prior to the increase in *p*, which leads to the different on-to-off and off-to-on paths.

**FIG. 3.**
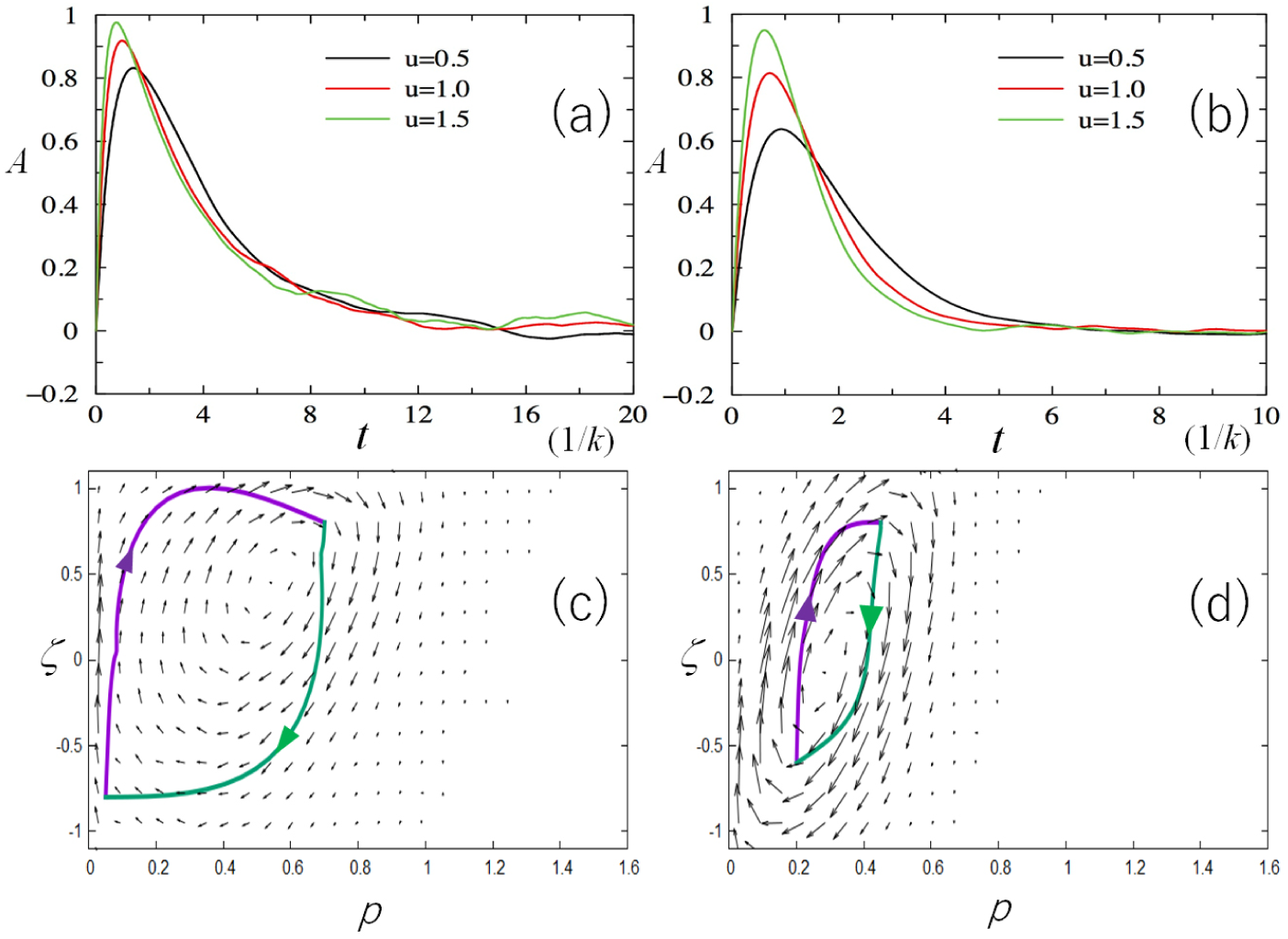
Temporal correlation and optimal paths in self-regulating genes. (a, c) Self-activating and (b, d) self-repressing genes. (a, b) The normalized difference between two-time cross correlations, *A*(*t*), is plotted as a function of *t* in units of 1*/k*, where *A*(*t*) > 0 represents the tendency that the change in the histone state affects the change in the amount of TF at time *t*. (c, d) The optimal paths for the on-to-off (green) and off-to-on (purple) directions are superposed on the flux **J** on the 2D plane of the protein concentration *p* and the histone state *ζ*. Calculated using the adiabatic approximation of TF binding/unbinding. The rates of the histone state change are scaled by the parameter *u* as *r*_10_ = *r*_0−1_ = *r*_01_ = *r*_−10_ = *u* with Δ*r*_10_ = Δ*r*_0−1_ = *u* in (a, c) or Δ*r*_01_ = Δ*r*_−10_ = *u* in (b, d). In (a) and (b), *A*(*t*) is plotted with *u* = 1.5 (green), *u* = 1 (red), and *u* = 0.5 (black). In (c) and (d), *u* = 1. The other parameters of (a, c) are same as in Fig. 1b and those of (b, d) are same as in Fig. 1d.

The hysteresis of switching dynamics is shown by calculating the optimal paths of transitions. An optimal path is obtained by first setting the start and end points of the path and then minimizing the effective action appearing in the path-integral formalism of kinetics connecting those points [3, 19, 33, 36–39]. Figs. 3c and 3d show the paths calculated by the simulated annealing optimization of the action using the algorithm in [3]. Thus obtained two paths, off-to-on and on-to-off paths, are indeed different from each other, both of which are consistent in their route orientations with the circulating probability flux. Because equilibrium kinetic paths should pass through the same saddle point of the landscape in both directions without showing the hysteresis, the calculated hysteresis is a manifestation of the nonequilibrium feature of gene switching. Furthermore, the area formed by the loop of forward and backward paths gives a quantitative measure of the heat dissipation and is linked to the nonequilibrium geometric phase [40].

When the TF binding/unbinding is slow, we need to solve Eq. 3 without using the adiabatic approximation. Slow non-adiabatic binding/unbinding were quantitatively examined by tuning the binding rate in bacterial cells in the recent experiments [41, 42]. Fig. 4a shows the landscape and flux in such a non-adiabatic case with the intermediate binding affinity of the activator TF. We find two basins for the on and off states and a distinct circulating flux between them. The qualitative feature is same as in the adiabatic case; however, here, we find the correlated binding/unbinding behavior with the histone state change, which should generate hysteresis in the 3D space. This hysteresis can be found in the calculated optimal paths in the 3D space (Fig. 4b).

**FIG. 4.**
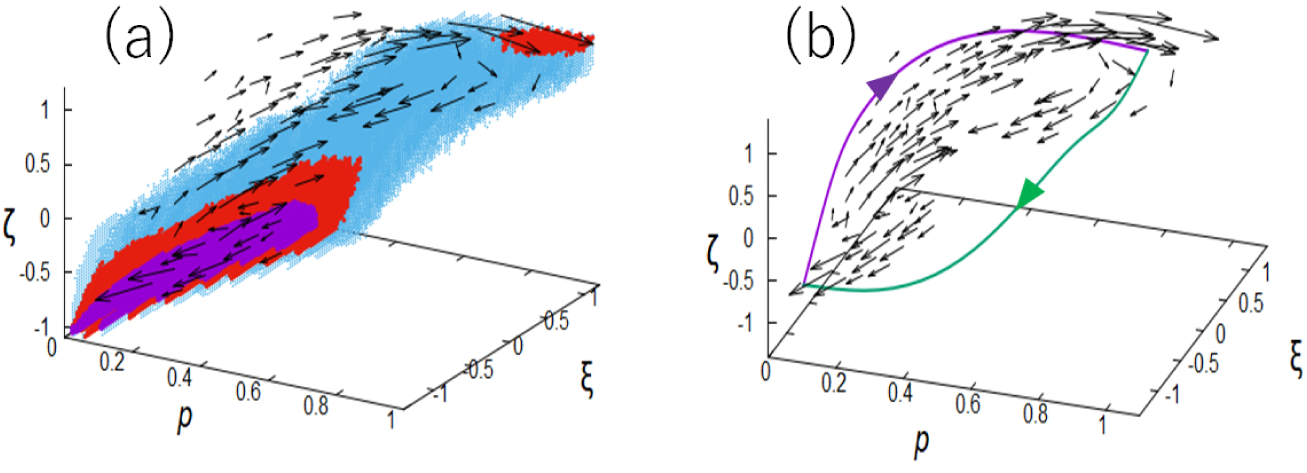
Landscape, probability flux, and optimal paths of a self-activating gene calculated in the 3D space of the protein concentration *p*, the TF binding status *ξ*, and the histone state *ζ*, in the non-adiabatic kinetics. (a) Flux **J** is superposed on the landscape *U* with 1 < *U* ≤ 2 (light blue), –0.5 < *U* ≤ 1 (red) and *U* ≤ *–*0.5 (purple). We find a gene-off state at *p* ≈ 0, *ξ* ≈ – 1, and *ζ* ≈ 0 ∼ –1 and a gene-on state at *p* ≈ *ξ* ≈ *ζ* ≈ 1. (b) Flux and optimal paths of on-to-off (green) and off-to-on (purple) directions. The rate parameters are scaled in units of *k* with *h*_0_ = 2 and *f* = 0.1. The other parameters are same as in Fig. 1b.

The flux-landscape approach provides a clue to resolving the ‘chicken-and-egg’ argument on the causality between the histone state and TF binding. The landscape analyses showed that the stability of each histone state is determined by the binding affinity of TF, which is in accord with the general belief that the specific TF binding is the cause and the histone state change is the result [43]. However, unexpectedly, the histone state tends to change in self-regulating gene circuits prior to the change in the amount of TF, which induces hysteresis in the switching dynamics; the histone state fluctuation can be a trigger for switching the feedback loop. This predicted temporal correlation needs to be tested by statistically analyzing *A*(*t*) in the single-cell experimental data.

Finally, the flux-landscape method should be applicable to problems of other epigenetic degrees of freedom. For example, a Monte Carlo type simulation of a gene network suggested that formation/dissolution of a super-enhancer of Nanog induces a large fluctuation in the gene activity in ESCs [7]. It is intriguing to develop a method to explain the degree of freedom of super-enhancer formation/dissolution and the associating large-scale chromatin structural change by extending the present scheme. Thus, the flux-landscape approach should provide insights in the problems of stochastic biological switching.

B. B. and M. S. would like to thank Dr. S. S. Ashwin for fruitful discussions. This work was supported by JST-CREST Grant JPMJCR15G2, the Riken Pioneering Project, and JSPS-KAKENHI Grants JP19H01860 and 19H05258. J. W. thanks NSF-PHY-76066 for supports.

